# iLEC-DNA: Identifying Long Extra-chromosomal Circular DNA by Fusing Sequence-derived Features of Physicochemical Properties and Nucleotide Distribution Patterns

**DOI:** 10.1101/2023.09.01.555875

**Authors:** Ahtisham Fazeel Abbasi, Muhammad Nabeel Asim, Andreas Dengel, Sheraz Ahmed

**Affiliations:** Rhineland-Palatinate Technical University of Kaiserslautern-Landau, Department of Computer Science, Kaiserslautern, 67663, Germany; German Research Center for Artificial Intelligence GmbH, Kaiserslautern, 67663, Germany

## Abstract

Long extrachromosomal circular DNA (leccDNA) regulates several biological processes such as genomic instability, gene amplification, and oncogenesis. The identification of leccDNA holds significant importance to investigate its potential associations with cancer, autoimmune, cardiovascular, and neurological diseases. In addition, understanding these associations can provide valuable insights about disease mechanisms and potential therapeutic approaches. Conventionally, wet lab-based methods are utilized to identify leccDNA, which are hindered by the need for prior knowledge, and resource-intensive processes, potentially limiting their broader applicability. To empower the process of leccDNA identification across multiple species, the paper in hand presents the very first computational predictor. The proposed iLEC-DNA predictor makes use of SVM classifier along with sequence-derived nucleotide distribution patterns and physicochemical properties-based features. In addition, the study introduces a set of 12 benchmark leccDNA datasets related to three species, namely HM, AT, and YS. It performs large-scale experimentation across 12 benchmark datasets under different experimental settings using the proposed predictor and more than 140 baseline predictors. The proposed predictor outperforms baseline predictors across diverse leccDNA datasets by producing average performance values of 80.699%, 61.45% and 80.7% in terms of ACC, MCC and AUC-ROC across all the datasets. The source code of the proposed and baseline predictors is available at https://github.com/FAhtisham/Extrachrosmosomal-DNA-Prediction.

## Introduction

Deoxyribonucleic acid (DNA) is comprised of billions of nucleotides and special arrangements of these nucleotides contain essential information for the development, functioning and inheritance of living organisms^1,2^. These nucleotides represent 25,000 protein-coding genes and various regulatory elements that control gene regulation^2^. In DNA sequence, these genetic components are organized in a structured manner and the sequence is wrapped around histone octamers also known as nucleosomes. Together around 30 million nucleosomes lead to the formation of chromosomes^3^. These chromosomes control vital biological processes like gene regulation, DNA replication, DNA damage response, and cell division^1,4^. However, aberrations within these processes produce additional genetic elements like extrachromosomal circular DNA (eccDNA)^5^.

To grasp the concept of eccDNA formation, one can examine the process of cell division^5,6^. During cell division, DNA replicates itself to ensure the transmission of chromosomes from parent to child cell. Within the replication process, DNA can incur damages, subsequently resulting in the fragmentation of chromosomes. DNA repair mechanisms reassemble these smaller segments and during the reassembling process, apart from chromosomes as a by-product eccDNAs are produced^5,6^. The lengths of these eccDNAs range from a few hundred to several thousand nucleotides because they are generated through random combinations of multiple segments^6^. Such fragments often harbor protein-coding genes, further complicating their impact on cellular processes such as gene expression, DNA replication, and DNA damage response.

EccDNAs can be classified into two distinct categories based on their size and characteristics: short eccDNAs and long eccDNAs (leccDNA). EccDNAs with shorter lengths typically have tens to a few hundred nucleotides^6,7^. They are commonly found in the nucleus and cytoplasm of the cell and facilitate movement between genomic loci and drive genetic diversity along with adaptation. Moreover, they store genetic information and replicate independently as episomes. On the other hand, leccDNAs are longer and contain thousands of nucleotides^8^. They are found only in the nucleus and contribute to genomic instability, gene amplification, cellular adaptation, and gene expression^6^. The presence of both types of eccDNA leads to excessive production of specific proteins, including oncogenes, enhancing the cell oncogenic potential and driving uncontrolled cell growth^9^. Futher, eccDNAs contribute to various diseases in multiple systems, such as glioblastoma, neuroblastoma, irregular immune response, and myocardial infarction^6,10–12^.

Identification of short eccDNA can reveal their roles in gene transfer, and genetic diversity. It is useful in understanding the molecular events of oncogene over-expression and therapeutic resistance. LeccDNA identification provides useful information about indications of genomic instability, genome organization as well as gene regulation. Furthermore, its identification is also useful for unveiling potential mechanisms responsible for the initiation and propagation of diseases such as cancer. Researchers are actively trying to explore its potential as a cancer biomarker and therapeutic resistance indicator^7,13^.

The identification of eccDNA is accomplished using a variety of wet-lab experimental methods, including pulsed-field gel electrophoresis (PFGE)^14^, southern blotting^15^, whole genome sequencing (WGS)^16^, fluorescence in situ hybridization (FISH)^17^, RT-PCR^15^, electron microscopy^18^, and rolling circle amplification (RCA)^19^. However, these methods often require prior knowledge or specific probes capable to bind with eccDNA which can limit their applicability to previously characterized eccDNAs. In addition, it is quite laborious, expensive, and time-consuming to identify eccDNAs at a larger scale across different organisms or cells.

The limitations of wet lab based methods and exceptional performance of AI based applications in natural language processing (NLP) tasks, have prompted a marathon of developing AI methods for DNA sequence analysis. Several AI models have been developed for various DNA analysis tasks such as enhancer identification^20,21^, DNA modification prediction^22,23^, promoter prediction^24^, DNA cyclizability prediction^25^, nucleosome position detection^26^ and so on. On the other hand, the identification of eccDNA is still being performed through wet lab-based methods due to the deficiency of AI applications for this particular task. According to the best of our knowledge, one predictor named DeepCircle^27^ is developed for the identification of short eccDNA sequences. There is currently no single predictor available for the identification of leccDNA, and also DeepCircle is not suitable for this specific purpose. The primary obstacle in utilizing DeepCircle for leccDNA identification lies in its reliance on the BERT model^27^, which can only handle sequence lengths of up to 512 tokens and leccDNA sequences exceed this token limit^28^.

In order to expedite and enhance research pertaining to the identification of leccDNA, there is an urgent necessity of a robust computational predictor. With an aim to develop a robust and precise computational predictor for leccDNA identification, the contributions of this study are manifold. Following the need for leccDNA identification datasets, it presents 12 benchmark datasets related to leccDNA sequences belonging to 3 different species i.e., *Homo sapiens* (HM), *Arabidopsis thaliana* (AT), and *Saccharomyces cerevisiae* (SC). It presents a robust and precise iLEC-DNA predictor that reaps the benefits of 2 different encoding methods for transforming raw sequences into statistical vectors. Furthermore, to discriminate leccDNA and non-leccDNA sequences, it employs SVM classifier that extracts more useful discriminative features from statistical vectors having nucleotide distribution patterns and physiochemical properties based information. Furthermore, it compares the performance of proposed predictor with more than 140 baseline predictors that are developed by using 13 most widely used sequence encoding methods and 11 machine learning classifiers. It conducts extensive experimentation over 3 different species datasets to find important answers to the following research questions; I) Do leccDNA sequences exhibit any distinctive nucleotide patterns that distinguish them from non-leccDNA sequences? II) How can variable-length leccDNA sequences be effectively handled to train ML classifiers? III) Which sequence encoding method is more competent in transforming raw leccDNA sequences into statistical vectors by incorporating discriminatory information? IV) Which sequence encoding method demonstrates better performance with which ML classifier? V) Does the combined potential of the multiple sequence encoding methods enhance the classification efficacy? VI) Does the ensembling of ML classifiers trained on the combination of different statistical vectors provide any classification improvement? We believe answers to these questions will provide valuable guidance to the research community when it comes to selecting the optimal combination of encoding methods and classifiers. This will significantly contribute to the creation of an efficient end-to-end predictive pipeline.

## Results

### Key Idea

Over the newly developed 12 benchmark leccDNA datasets, we generate an effective statistical representation based on the gap-kmer distribution and physiochemical properties of nucleotides using two sequence encoding methods namely, CKSNAP, and SCPSEDNC. Using discriminatory features from CKSNAP and SCPSEDNC we develop a novel predictor based on support vector machine (SVM) for leccDNA identification namely, iLEC-DNA. In order to prepare fixed-length leccDNA and non-leccDNA sequences without losing information-rich regions, we perform a thorough intrinsic 2-mer distribution analysis. To validate the observations from the intrinsic analyses, an extrinsic performance analysis is conducted which affirms the information-rich regions that play a critical role in leccDNA identification. In addition, we compared the performance of the proposed predictor with more than 140 baseline predictive pipelines developed based on 13 commonly used sequence encoding methods and 11 ML classifiers. Extensive experimentation shows that the proposed predictor is able to achieve suitable performance across diverse benchmark datasets for leccDNA identification.

### Summary of Results

This section provides a comprehensive overview of the research objectives pertaining to the prediction of leccDNA. First, it investigates whether leccDNA sequences exhibit nucleotides distinctive patterns that can differentiate them from non-leccDNA sequences. It illustrates the performance values of 143 baseline predictors across 5 different sequence lengths. It compares the performance of proposed and baseline predictors. Finally, it illustrates the performance values of the proposed leccDNA predictor on 12 benchmark datasets.

### Do leccDNA sequences exhibit any distinctive nucleotide patterns that distinguish them from non-leccDNA sequences?

In order to perform DNA sequence classification, ML predictors require uniform length of DNA sequences and distinct nucleotide patterns across various classes. As depicted in Figure 10, there is considerable variability in the lengths of both leccDNA and non-leccDNA sequences. However, for the purpose of training ML classifiers, these sequences must be of a fixed length. To tackle this issue, one solution involves the direct addition of a padding character ‘P’ within sequences. However, the substantial variations in leccDNA sequence lengths, spanning from five to thirty thousand nucleotides necessitate more padding values which introduce noise and bias in data. This influx of padding values not only disrupts the original data distribution but also undermines the model’s potential to generalize effectively on unseen data. In an alternate strategy^29^, first information-rich regions are explored in the sequences, and padding values are added after truncating all sequences with a specific length threshold. This approach produces fixed-length DNA sequences without introducing substantial bias in data.

With the objective of delineating information-rich regions and obtaining uniform-length DNA sequences, an initial step involves the segmentation of DNA sequences into discrete sub-sequences. We explored information-rich regions by generating 10 different types of subsequences including 0-500, 0-1000, 0-1500, 0-2000, 0-2500, 0-3000, 0-3500, 0-4000, 0-4500, and 0-5000 nucleotides. The rationale behind this range selection stems from the distribution analysis of sequence lengths, which manifests a notable skew towards lengths below 5000 nucleotides as shown in Figure 10.

To gain insights into the density and patterns of distinct nucleotide pairs, in 4 different steps an intrinsic analysis is performed. First, the occurrences of 16 unique 2-mers within each sub-sequence are calculated in leccDNA and non-leccDNA sub-sequences. Next, The sub-sequence-based densities of 2-mers are computed separately for leccDNA and non-leccDNA sequences across 10 different sequence lengths. In the subsequent step, the density-based values are normalized with a total number of sequences. Finally, the density differences of 2-mers among leccDNA and non-leccDNA sequences are computed to reveal distinctive nucleotide patterns.

Figure 1 shows the sub-sequence density-based differences of different 2-mers across 3 different leccDNA benchmark datasets namely, C4-2B, AT, and YS. The majority of density-based differences lie within the initial sub-sequences, whereas among the last sub-sequences, the densities of 2-mers are similar among leccDNA and non-leccDNA sequences. These differences in the densities of specific nucleotide pairs indicate that certain regions of the DNA sequences exhibit distinct nucleotide distribution patterns, which are characteristic of leccDNA sequences and can differentiate leccDNA from non-leccDNA sequences. Particularly certain 2-mers such as TT, TC, TA, AT, AC, GG, GC, and GA, have notably higher densities in leccDNA sequences, and 2-mers i.e., AG, GC, TG, CA, GA, and TC, show higher densities in non-leccDNA sequences. These distinguishing factors are captured with the help of specific sequence encoding methods and can be utilized for the identification of leccDNA sequences. In addition, similar patterns and 2-mer density-based differences across leccDNA and non-leccDNA sequences of other benchmark datasets are provided in supplementary file x.

**Figure 1.**
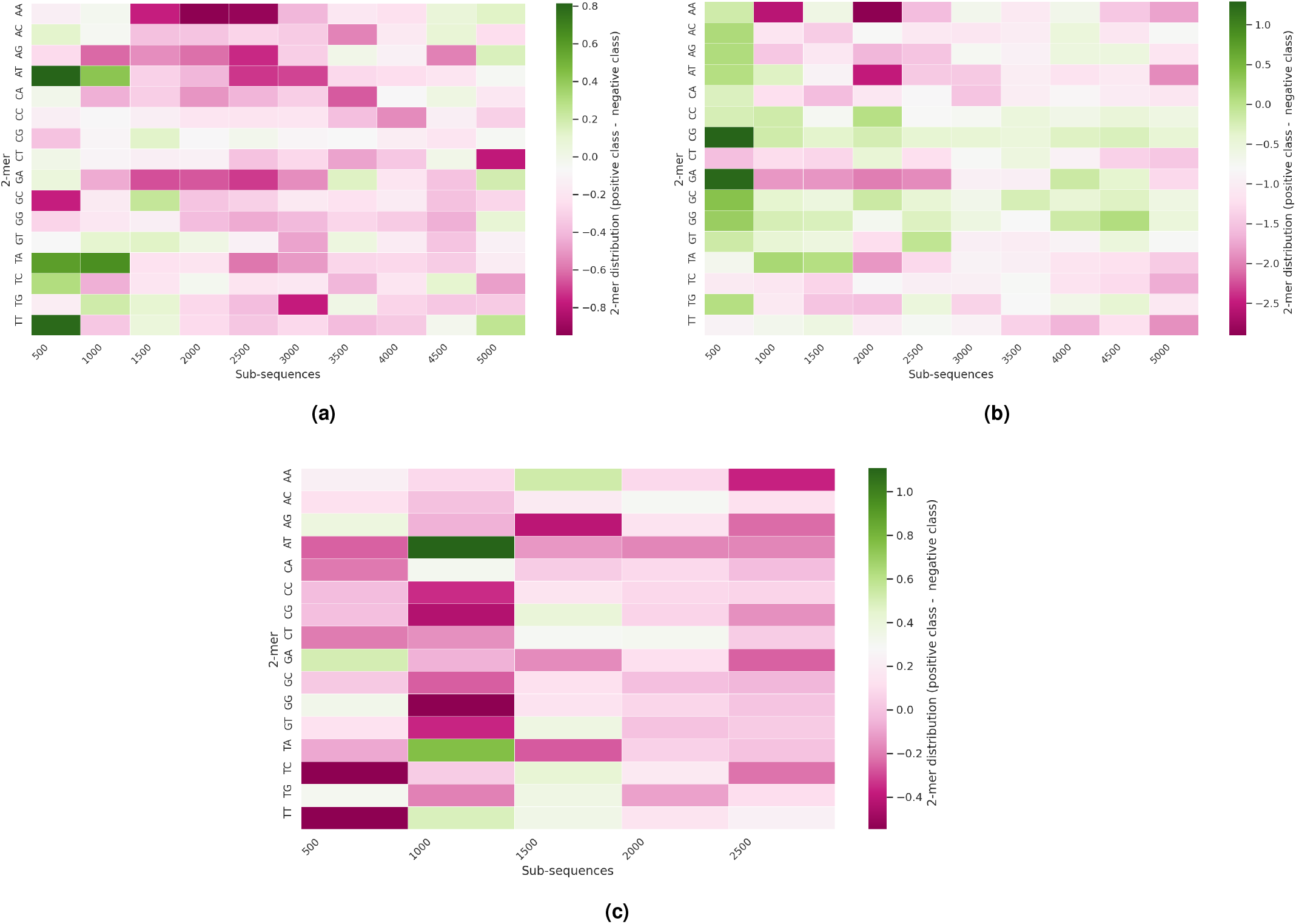
Sub-sequence-based density distribution of k-mers in 3 different benchmark datasets i.e. (a) C4-2B, (b) AT, and (c) YS.

The intrinsic nucleotide pattern analysis affirms that leccDNA sequences display discernible nucleotide patterns that differentiate them from non-leccDNA sequences. These distinguishing features are predominantly situated in the initial regions of leccDNA sequences which are further investigated in the subsequent extrinsic performance analyses, as elaborated in the following subsection.

### How can variable length leccDNA sequences be effectively handled to train ML classifiers?

With an aim to analyze the impact of 5 different regions (1000, 2000, 3000, 4000, and 5000) in leccDNA identification, using 13 encoding methods and 11 classifiers based 143 predictive pipelines an extrinsic performance outcome is discussed in this section.

Figure 2 illustrates top performing baseline predictors ACC values, across 5 different sequence lengths for 12 distinct benchmark datasets under independent test setting. Two different types of performance trends are observed across benchmark leccDNA datasets with respect to sequence lengths i.e., I) steady linear performance increase with sequence length, II) linear performance increase up to a specific sequence length. The leccDNA benchmark datasets namely, C4-2B, FAD, and SP lie under the trend category I as they have a gradual increase in performance with respect to the increase in the sequence length. These benchmark datasets have maximum performance value at the sequence length of 5000. In addition, the rest of the benchmark datasets fall under the trend category II, as the performance values increase up to a certain sequence length and afterward the performance decreases. For instance, OVCAR8, YS, PC, and FADU show a gradual increase in performance up to the 3000 nucleotides and a performance decline afterward. In addition, OP, UR, and AT show similar patterns until 2000 and PL, and MS show performance improvement up to 4000 nucleotides after which the performance deteriorates.

**Figure 2.**
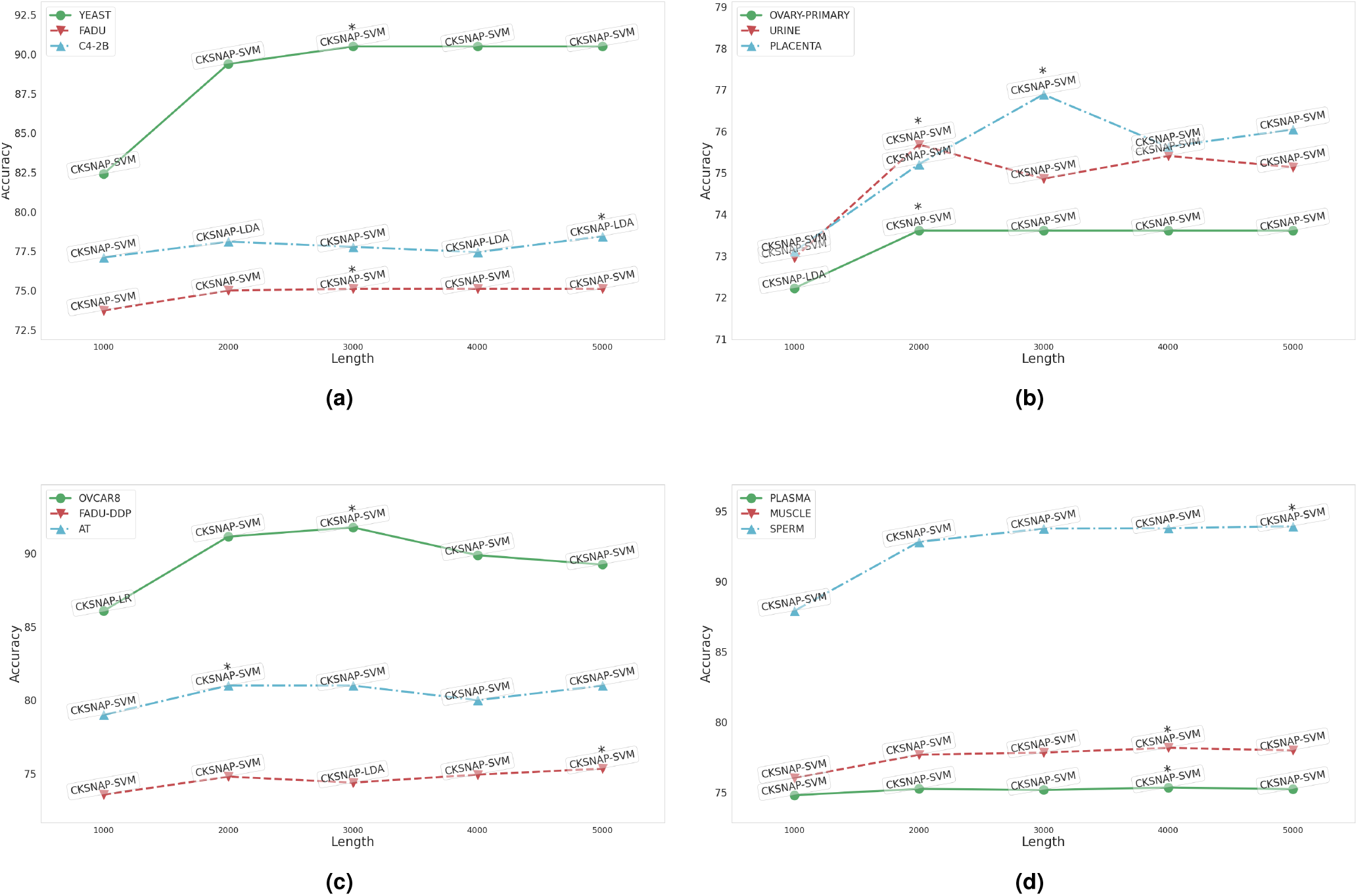
Performance comparison of 13 sequence encoding methods and 11 ML classifiers in terms of ACC over 12 leccDNA benchmark datasets at 5 different sequence lengths: 1000, 2000, 3000, 4000, and 5000.

It is important to note that while dealing with leccDNA sequences, extracting meaningful information from subsequences that contain initial sequence nucleotides proves to be advantageous in achieving optimal classification efficacy. This highlights the potential benefits of breaking down complex sequences into manageable chunks, allowing for better classification performance and more efficient analysis. Furthermore, the superior performance of the baseline predictors for the initial sequence lengths reinforces the previously discussed observations that most of the discriminative patterns related to nucleotides are concentrated in the initial regions of leccDNA sequences.

### Which sequence encoding method demonstrates better performance with which ML classifier?

This section briefly addresses research questions (III and IV) pertaining to the optimal sequence encoding methods and ML classifiers for effective leccDNA identification. To achieve this, two analyses are conducted here, first the performance rank scores of each sequence encoding method are calculated across diverse classifiers on 7 datasets. Additionally, the rank scores of classifiers are computed to identify consistently superior classifiers across all sequence encoding methods. Figure 3a shows the rank scores of 13 different sequence encoding methods with 11 ML classifiers across 7 datasets namely, AT, C4-2B, OP, OV, PL, UR, and YS. The rank scores are computed by determining the maximum performance of a sequence encoding method across a classifier for different datasets.

**Figure 3.**
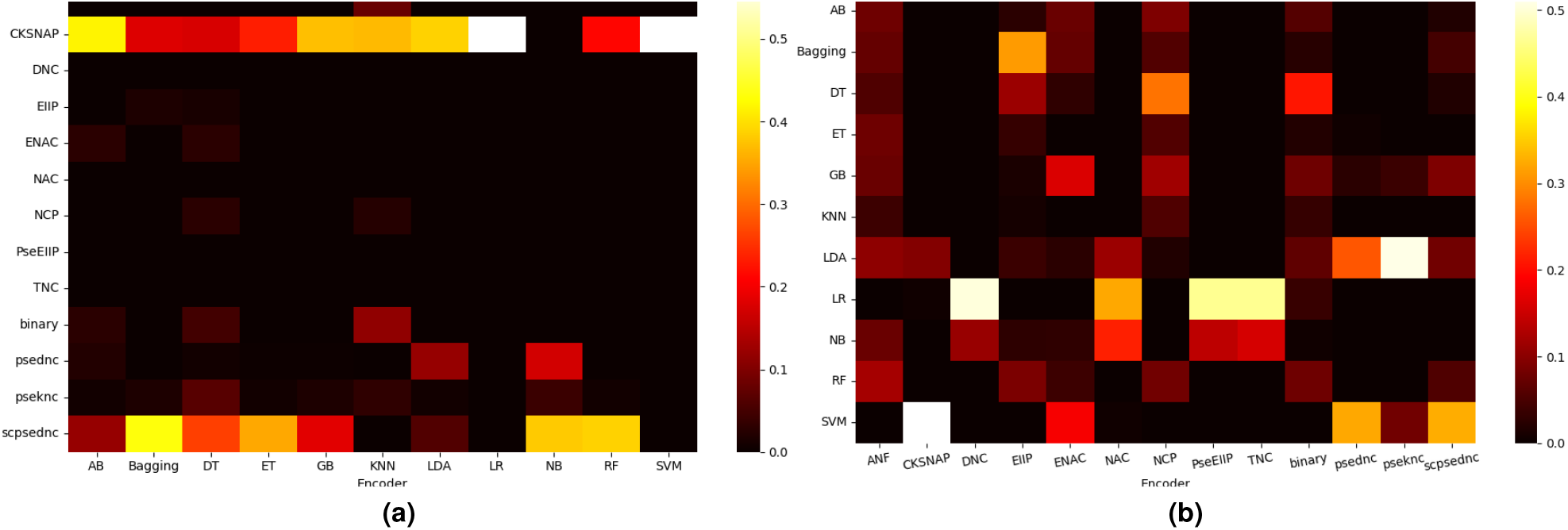
(a) Unraveling the top-ranked scores of 13 sequence encoding methods across 11 classifiers over multiple datasets (AT, C4-2B, OP, OV, PL, UR, and YS) and sequence lengths: 1000, 2000, 3000, 4000, and 5000. (b) Exploring the ranking scores of 11 ML classifiers on 13 sequence encoding methods, shedding light on their comparative performance.

Three distinct categories of sequence encoders are established based on performance rank scores: I) encoders with the lowest rank scores, II) encoders with inconsistent performance across certain classifiers, III) encoders with consistent performance across all classifiers. Among 13 sequence encoding methods, DNC, NAC, TNC, and EIIP fall under category I as these methods consistently show the lowest rank scores across 11 different ML classifiers. This attributes to the limited discriminatory power of these methods in capturing relevant patterns, nuanced variations, and characteristics in leccDNA sequences. Similarly, ANF, ENAC, NCP, PSEDNC and binary fall under category II as these methods show consistent ranks scores across few ML classifiers such as KNN, AB, DT, NB and LDA. Particularly, two sequence encoders namely, CKSNAP and SCPSEDNC lie under category III as they show consistent rank scores across majority of the classifiers. The consistent performance of these methods lies with their ability to efficiently capture nucleotide distribution and physiochemical properties (base stacking, and stability in terms of SCPSEDNC) that enable ML classifiers to identify leccDNA sequences with more efficacy as compared to the other sequence encoding methods.

Figure 3b shows the rank scores of 11 different ML classifiers with 13 distinct sequence encoding methods across AT, C4-2B, OP, OV, PL, UR, and YS datasets. Notably, the KNN classifier consistently exhibits the lowest predictive performance across all the sequence encoding methods. Similarly, the AB classifier demonstrates inconsistent and comparatively lower performance ranks across all sequence encoding methods. Various classifiers including GB, bagging, ET, DT, and NB exhibit relatively inconsistent performance rank scores, as they have marginal rank scores only with EIIP, NCP, and binary sequence encoding methods. On the other hand, SVM, LDA, RF and LR classifiers stand out with the highest rank scores, showing their effectiveness in the identification of leccDNA sequences with diverse sequence encoding methods.

These analyses confirm that among diverse sequence encoding methods two methods, CKSNAP and SCPSEDNC encode more discriminatory patterns in statistical vectors that help ML classifiers in leccDNA identification. Overall, among 11 classifiers only SVM, LDA, RF, and LR produce robust performance and are well-suited for accurate leccDNA identification.

### Does the combined potential of the multiple sequence encoding methods enhance the classification efficacy?

To find the answer to research question V, we explore the potential of 2 top-performing sequence encoding methods (CKSNAP and SCPSEDNC) and 3 top-performing classifiers (SVM, LDA, RF). Here the objective is to analyze whether these 3 classifiers produce better performance with statistical vectors of standalone encoders or with their combined statistical vectors. Furthermore, as discussed earlier, among 13 sequence encoders and 11 classifiers, two encoders (CKSNAP, SCPSEDNC) and 3 classifiers produced the best performance. Here among 143 baseline predictive pipelines, we performed a performance comparison of top performing 6 predictive pipelines with proposed predictor that makes use of combined the potential of two encoders (CKSNAP, SCPSEDNC) and SVM classifier. Moreover, in addition to 6 top performing baseline predictors, we compare the proposed predictor performance with two special predictive pipelines that also utilize the combined potential of two encoders (CKSNAP, SCPSEDNC) with LDA and RF classifiers. Figure 4 showcases the evaluation results based on performance values generated by top performing baseline and proposed predictors. It highlights the performance gains achieved through the utilization of diverse discriminatory features from CKSNAP and SCPSEDNC across other ML classifiers such as LDA and RF.

**Figure 4.**
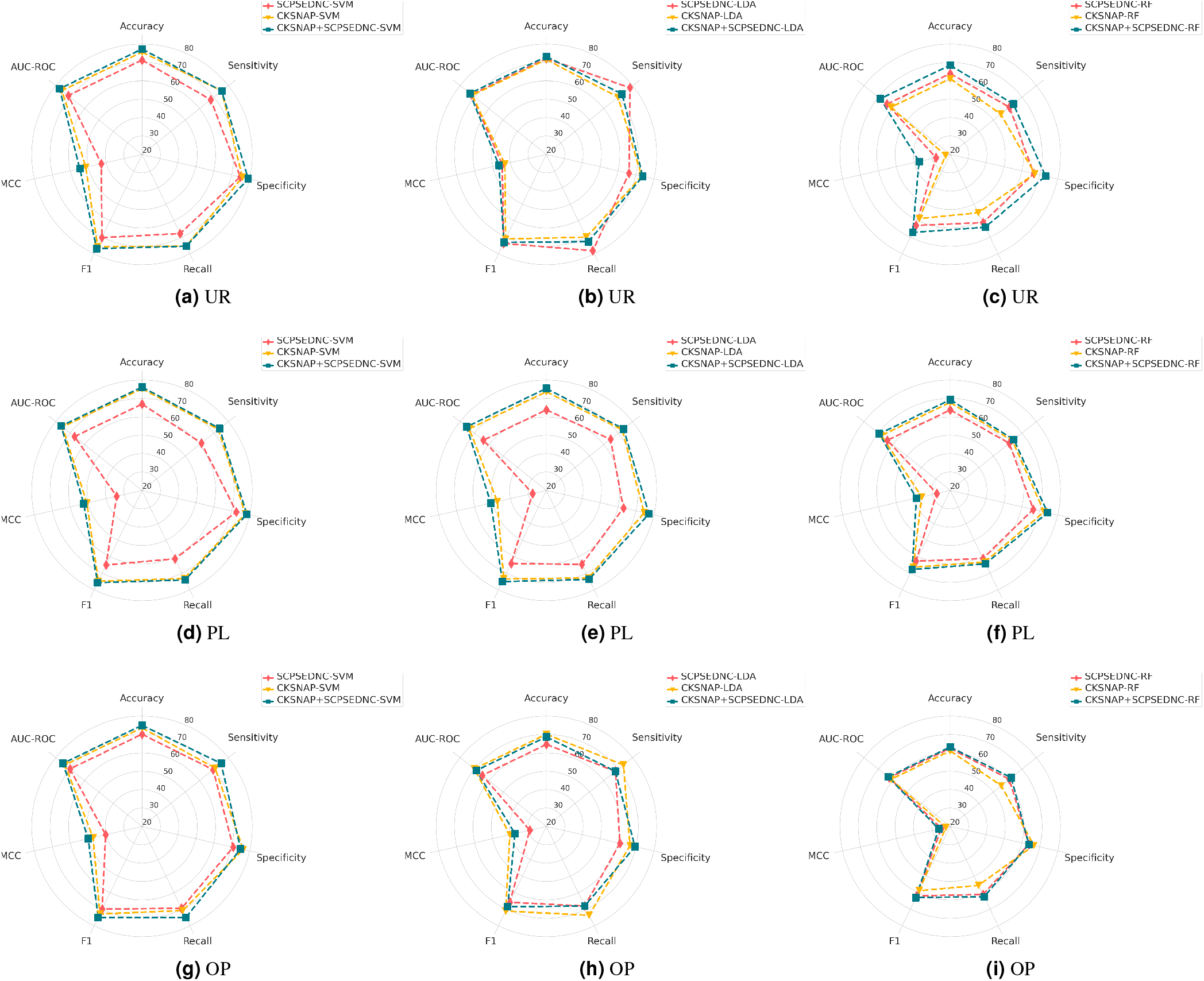
Performance scores of 3 different datasets over 1st and second stage of classification. Each row represents a unqiue dataset i.e., UR, PL, and OP, whereas, (a) shows the performance scores of SVM, (b) demonstrates the performance scores for LDA, and (c) represents the performance scores for RF classifier over 6 distinct evaluation measures namely, ACC, SN, SP, MCC, F1, and AUC-ROC.

In terms of UR dataset, the collective potential of CKSNAP and SCPSCDNC demonstrates an average performance enhancement of 3.824% with SVM, 1.366% across LDA, and a substantial increase of 6.011% across RF with respect to the ACC values of top performing baseline predictive pipelines. Similarly, for PL dataset combined statistical vectors from CKSNAP and SCPSEDNC show an average performance gain of 5.07% with SVM, 6.772% with LDA and 3.505% with RF in terms of ACC. Futhermore, in terms of OP dataset the combined statistical vectors illustrate performance gains of 3.125% with SVM, 1.33% with LDA, and 1.391% with RF in terms of ACC as compared to top-performing baseline predictors.

Overall, it is observed that both sequence encoding methods provide unique and discriminatory information to the classifiers for leccDNA identification. This discriminatory and unique information when presented to the ML classifiers in a concatenated way leads to signification performance gains which suggests the importance of using multiple sets of information while training a classifier for leccDNA identification. In addition, it is also observed that the proposed predictor produces better performance as compared to other predictors namely, LDA and RF. Therefore, the final experimentation over leccDNA benchmark datasets is performed by utilizing SVM, CKSNAP, and SCPSEDNC.

### Ensembling of Different Classifiers

With the primary goal of increasing the efficacy and robustness of the proposed predictive framework, the potential of ensembling for the identification of leccDNA sequences is also explored. Ensembling is carried out by using the probabilistic feature space of classifier outputs. Four top-performing classifiers, SVM, RF, LDA, and LR, are utilized for ensembling, that are initially trained on the combination of statistical vectors generated through CKSNAP and SCPSEDNC. The probabilistic outputs from these top-performing classifiers are combined in various ways and then passed through three additional classifiers, RF, LR, and SVM, for further classification. Despite conducting extensive performance analyses, no single combination of classifiers demonstrates consistent performance gains across all datasets. Nevertheless, the results for six distinct evaluation measures i.e., ACC, SP, SN, MCC, F1, and AUC-ROC, are provided in the supplementary files.

### Performance Analyses Over 5-fold Cross Validation

Figure 5 illustrates the performance results of the proposed leccDNA predictor in terms of 5-fold cross-validation across 12 benchmark leccDNA datasets. The proposed predictor shows high-performance values over the dataset of SP, OVCAR8, and YS ranging from 87-93% in terms of ACC, and AUC-ROC. For OVCAR8 and YS datasets there is an average gap of 1.9466% in terms of SP and SN, which suggests that the proposed predictor is not prone to type I and type II error. This implies that the proposed leccDNA predictor is quite robust in predicting samples belonging to positive and negative classes. The high performance of the proposed predictor is due to the sufficient number of samples present to train the proposed predictor.

**Figure 5.**
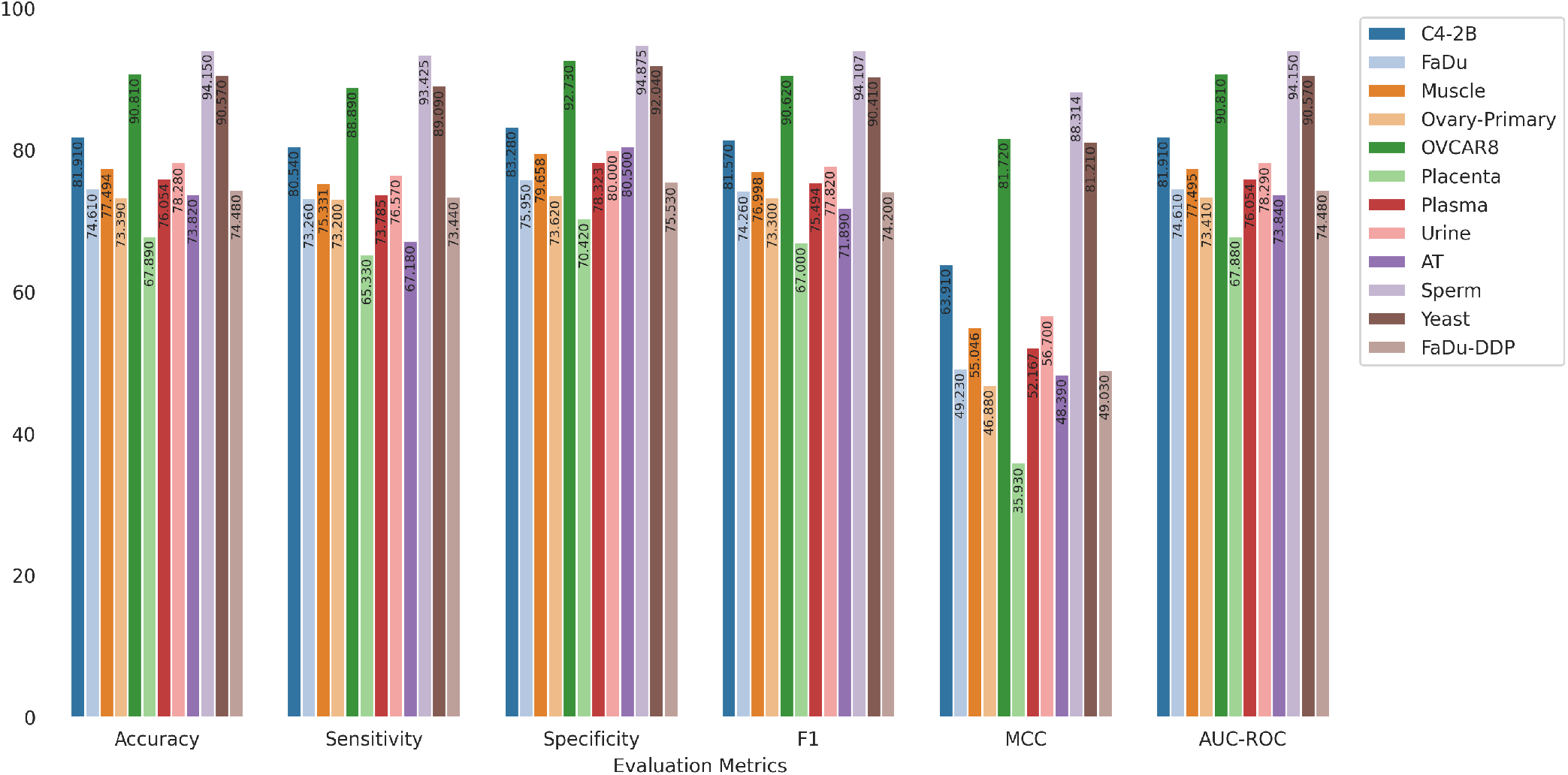
Performance values of the proposed predictor across 12 different leccDNA datasets over 6 distinct evaluation measures in terms of 5-fold validation.

In addition, across the datasets of OP, FaDu, PL, MS, UR, C4-2B, and FAD, the performance of the proposed predictor ranges from 73-80% in terms of ACC and AUC-ROC. The proposed predictor is not highly prone to type I and type II errors due to an average gap of 3.03% among SP and SN values. Moreover, the proposed predictor shows low performance on the PC and AT datasets with the performance values ranging from 67-73% in terms of ACC and AUC-ROC. In addition, over both of the datasets, the proposed predictor is prone to either type I or type II error due to an average gap of 9.105% in terms of SP and SN and lower AUC-ROC. The predictor is more prone to type II error as it is not able to successfully identify positive samples with a higher ratio as compared to the negative samples. The low performance of the proposed predictor on these datasets is due to the presence of a limited number of samples for positive and negative class samples (200 leccDNA sequences for AT and 400 leccDNA sequences in terms of PC).

### Performance Analyses Over Independent Test Set

Figure 6 illustrates the performance analyses over 6 distinct evaluation measures across 12 different benchmark datasets in terms of independent test sets. A closer look at the performance values at the 5-fold validation and independent test sets reveals that the performance of the proposed predictor either remains the same, increases, or decreases as compared to the 5-fold validation across various datasets. The performance of the proposed predictor remains the same across 6 different datasets such as FaDu, FAD, PL, MS, YS, and C4-2B. Similarly, there is a slight decrease in the performance of the proposed predictor over OVCAR8, and UR datasets, and an increase in the performance of 4 datasets namely, SP, AT, PC, and OP.

**Figure 6.**
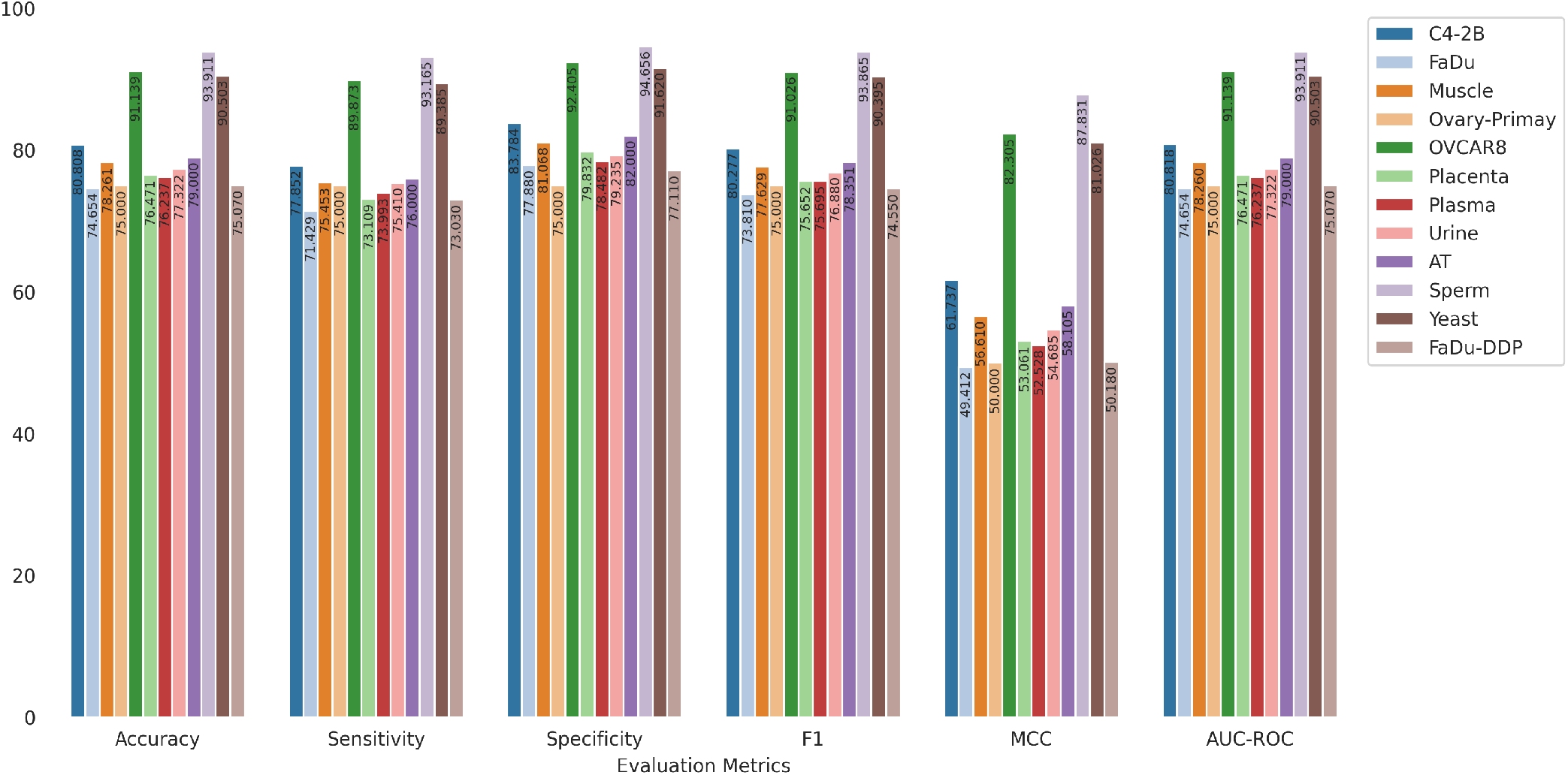
Performance values of the proposed predictor across 12 different leccDNA datasets over 6 distinct evaluation measures in terms of the independent test set.

## Discussion

Experimental results through intrinsic and extrinsic performance analyses reveal that initial regions of leccDNA carry siginifcant discriminatory information about nucleotide distribution for leccDNA identification. In addition, experimental results on 12 benchmark datasets from 3 different species, reveal that among 13 diverse types of encoding methods, two encoders CKSNAP and SCPSEDNC generate more comprehensive statistical vectors. A prime reason behind generating better statistical vectors is the extraction of both simple as well as gap-based nucleotide patterns. Specifically, CKSNAP encoder transforms raw DNA sequences into statistical vectors by computing occurrence distribution of simple as well as gap-based bimers. SCPSEDNC encoder also follows the footprints of CKSNAP, it makes use of precomputed physiochemical properties for computing nucleotides correlations at different positions. In a nutshell, it can be concluded, those encoding methods are more suitable for transforming raw DNA sequences into statistical vectors that emphasize on gap-based nucleotides distribution. Furthermore, concatenation of statistical vectors of both encoders facilitates the extraction of two different types of features, gap based bimers distribution and gap-based nucleotides correlational information extracted through physiochemical properties. Among 11 different classifiers, SVM remains top performer because it finds optimal hyperplanes for discriminating sequences into leccDNA and non-leccDNA classes. Experimental results reveal that across all species benchmark datasets, it successfully designed optimal hyperplanes. Although other classifiers produced better performance across different datasets but they are not well generalized such as apart from SVM classifier, RF, and LDA produced better performance but over 10 dataset LDA was the second top performer and over 2 datasets RF was second top performer.

Due to the substantial size of leccDNA sequences, a constrained set of sequence encoding methods is used to address computational inefficiencies and mitigate the increase in experimental count. Although the incorporation of distinct sequence order and nucleotide distribution information achieves a notable reduction in prediction errors, the robustness and efficacy of the proposed model are limited over the PL dataset due to a bias towards type I error. Moreover, in the future, we intend to leverage additional sequence encoding methods and incorporate certain deep learning models to enhance the classification efficacy and robustness across diverse leccDNA datasets.

## Conclusion

The primary objective of this study is to introduce a novel and effective predictive framework that can accurately predict leccDNA in various cell types and species. First, 12 different benchmark datasets are processed and formulated for leccDNA prediction. In addition, the unique distribution of nucleotides is explored with an aim to decode the discriminatory potential in leccDNA sequences. The proposed framework explores the potential of 13 different sequence encoding methods in conjunction with 11 ML classifiers to predict leccDNA. Comprehensive experiments reveal that SVM classifier and 2 sequence encoding methods namely, SCPSEDNC and CKSNAP give superior and consistent performance across diverse leccDNA datasets. We have observed that the concatenation of statistical vectors generated through CKSNAP and SCPSEDNC lead to significant performance gains. Based on various performance analysis, iLEC-DNA, a novel predictor for leccDNA, is proposed that captures the sequence order information through physiochemical properties such as stacking and stability, and nucleotide distribution information. iLEC-DNA is evaluated over 12 distinct benchmark datastes namely, MS, PS, SP, FaDU, FAD, PC, UR, CB, OV, OP, AT and YS. iLEC-DNA is a valuable tool for researchers examining intricate and lengthy eccDNA. Its capabilities can enable the exploration of leccDNA and their involvement in genomic instability and the onset of cancer.

## Materials and Methods

This section demonstrates comprehensive details of proposed and baseline predictors. It provides a comprehensive overview of benchmark datasets development process and characteristics of datasets. Finally, it presents evaluation measures that are used to evaluate and compare the performance of proposed and baseline predictors.

### Proposed iLEC-DNA Predictor

Figure 7 demonstrates different modules of the proposed iLEC-DNA predictor. It can be seen that after the datasets development process, DNA sequences are transformed into statistical vectors. The transformation of DNA sequences into statistical vectors is an essential task because AI predictors can only process numerical data and cannot operate directly on DNA sequences. While converting DNA sequences into statistical vectors the prime objective is to incorporate sequence order, semantic, nucleotide distribution, and positional features into the statistical vectors. The proposed predictor transforms raw DNA sequences into statistical vectors by reaping the combined benefits of two different types of sequence encoders namely, CKSNAP and SCPSEDNC. Finally, it concatenates the statistical vectors of both the encoders that is passed to the SVM classifier. A comprehensive description of both sequence encoding methods and their concatenation is provided following subsections.

**Figure 7.**
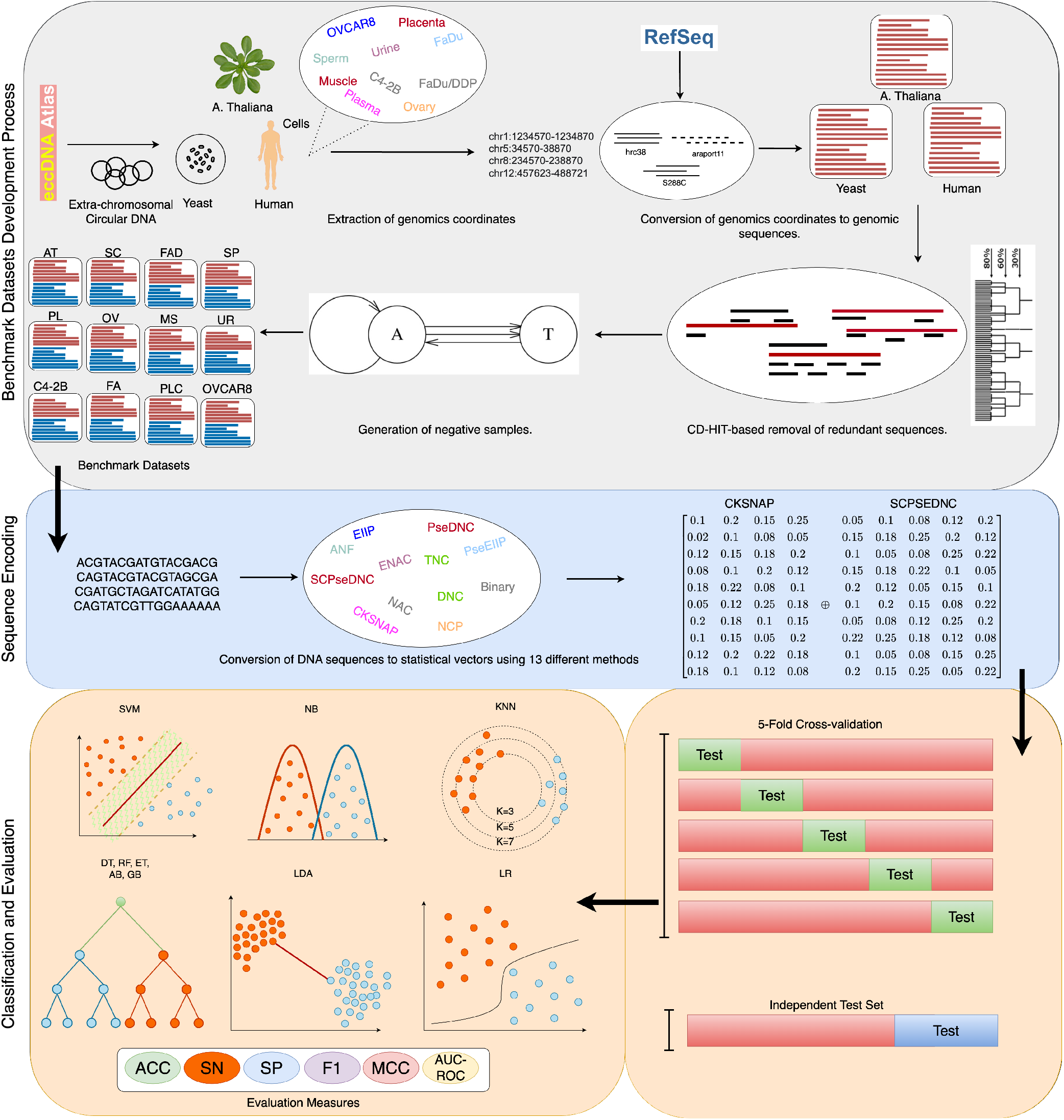
Graphical illustration of Benchmark datasets development process, proposed and baseline predictors. In datasets development process, leccDNA sequences are extracted from the eccDNA atlas and CD-HIT is utilized to remove redundant leccDNA sequences. Subsequently, USHUFFLE is applied to generate negative samples. In the second step, leccDNA sequences are converted into statistical vectors through baseline and proposed sequence encoding pipelines. In the classification and evaluation process, the performance of the proposed predictor is compared with the baseline predictor across all datasets.

#### Complementary K-spaced Nucleic Acid Pairs (CKSNAP)

CKSNAP encoder was proposed by Zhang et al.^30^ and has been widely used used in diverse types of DNA sequence classification predictors including, enhancer prediction^31^, DNA replication origin identification^32^, DNA modification prediction^33^ and promoter prediction^34^. The motivation behind the development of this encoder was to capture nucleotide occurrence distribution patterns with different gap values. CKSNAP^35^ generates gap based bimers such as for a hypothetical sequence GCTA, with gap value 1, gap-based bimers are be G-T and C-A. Similarly, for gap value 2, it first generates 1 gap-based bimers and then 2 gap-based bimers. Furthermore, for each gap value, it computes occurrence frequencies of bimers and normalizes them with total number of gap-kmers. Mathematically, CKNSAP can be written as;

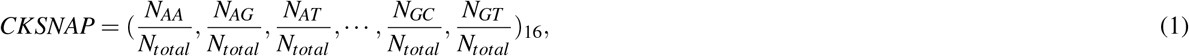

where, N_*AA*_ represents the total occurrences of bimer AA in the DNA sequence and N_*total*_ denotes the total number of gap bimers. A detailed working paradigm of CKSNAP is shown in Figure 8.

**Figure 8.**
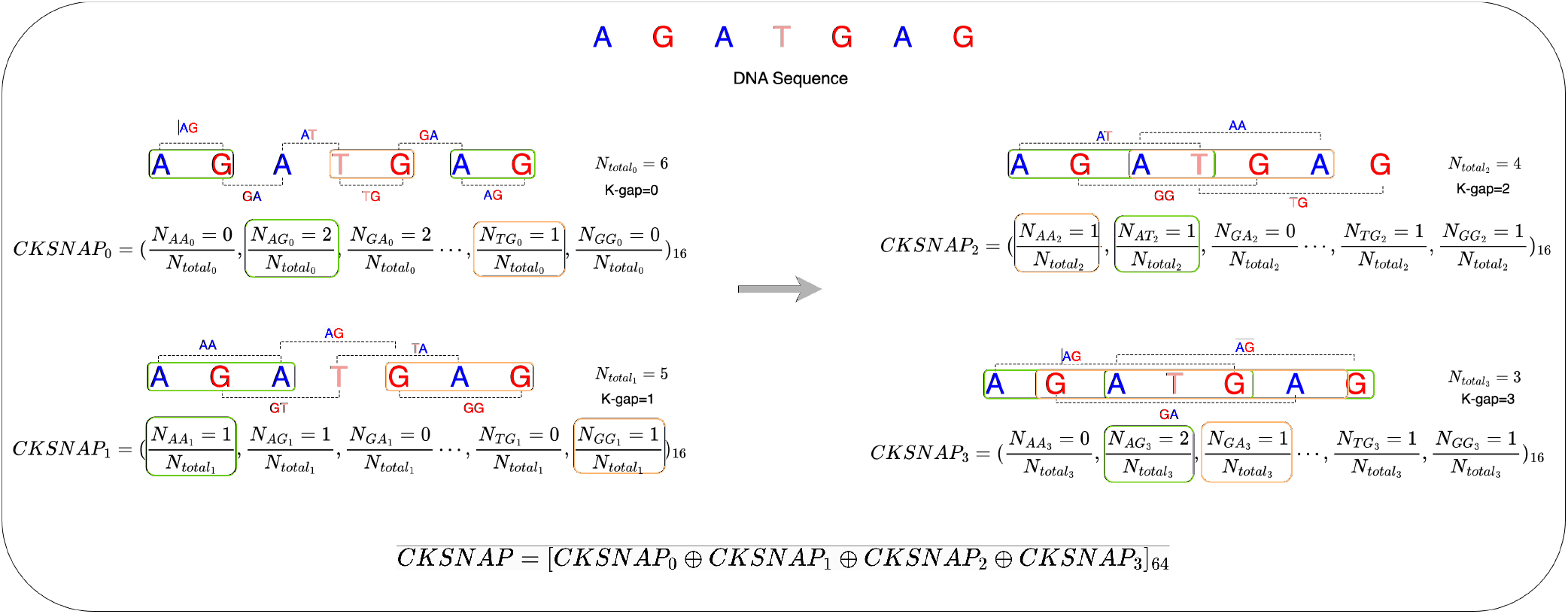
Working paradigm of CKSNAP sequence encoding method on a hypothetical DNA sequence i.e., AGATGAG with k-gap = 3.

#### Series Correlation Pseudo Dinucleotide Composition (SCPSEDNC)

SCPSEDNC encoder was proposed by Chen et al.^36^ and has been widely used in multiple DNA sequence classification applications including enhancer prediction^37^, DNA replication origin prediction^32^, and DNA hypersensitive site prediction^38^. This encoder transforms raw sequences into statistical vectors by computing two different types of information namely, nucleotide occurrence frequencies and correlation between nucleotides. To compute nucleotide occurrence frequencies, it generates bimers from the sequence and computes the occurrence frequencies of 16 unique bimers. Similarly, to compute correlational information of nucleotides, it computes the products of phsysiochemical indices among two different pairs of bimers that capture local and global patterns of nucleotides in DNA sequences. Mathematically, SCPSEDNC can be written as;

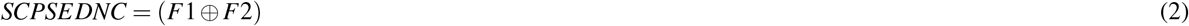

whereas, F1 can be formulated as,

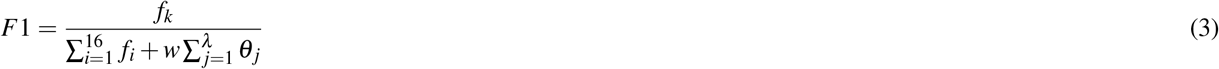

Here *f*_*k*_ represents the occurrences of a specific dinucleotide. 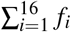 denotes the cumulative occurrences of all 16 dinucleotides in the DNA sequence. In addition, *F*_2_ can be written as,

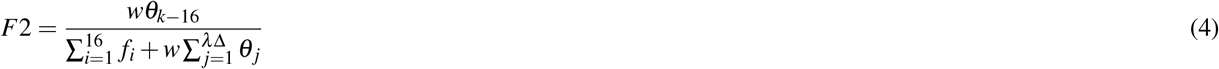

whereas, w is the weight term that ranges from 0 to 1. *λ* represents the highest counted rank of correlation and Δ denotes the number of physiochemical indices. *θ* can be written as;

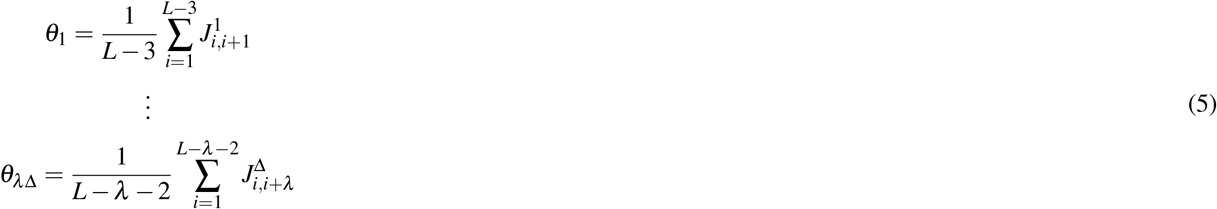

furthermore, 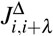 represents the correlation between two pairs of dinucleotides along a physiochemical index. A brief working paradigm of SCPSEDNC and selection of 2-mers with respect to lag value for correlation functions is presented in Figure 9.

**Figure 9.**
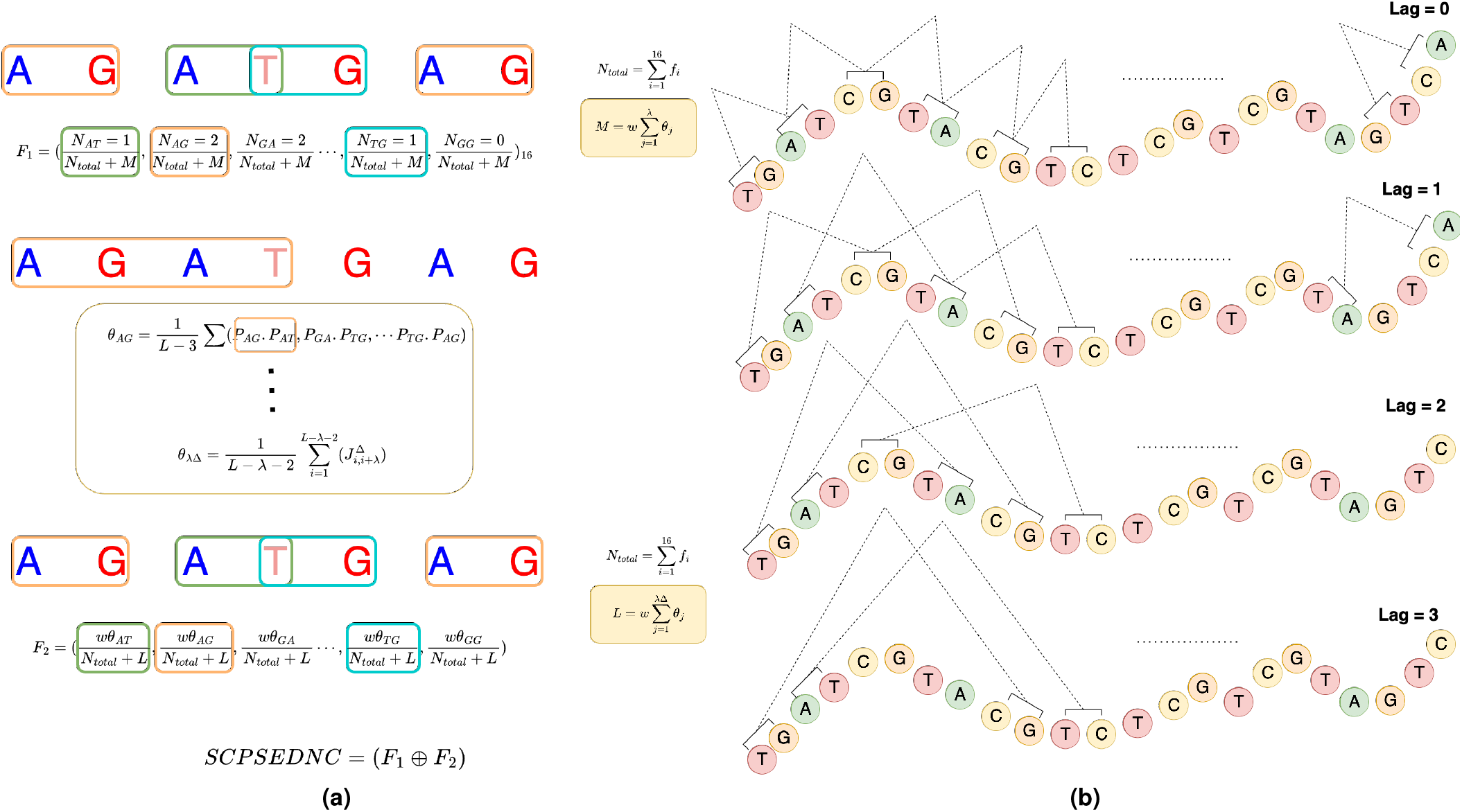
(a) Working paradigm of SCPSEDNC sequence encoding method on a hypothetical DNA sequence i.e., AGATGAG. (b) The selection of 2-mers with respect to a lag value for the computation of correlation functions on a hypothetical DNA sequence.

**Figure 10.**
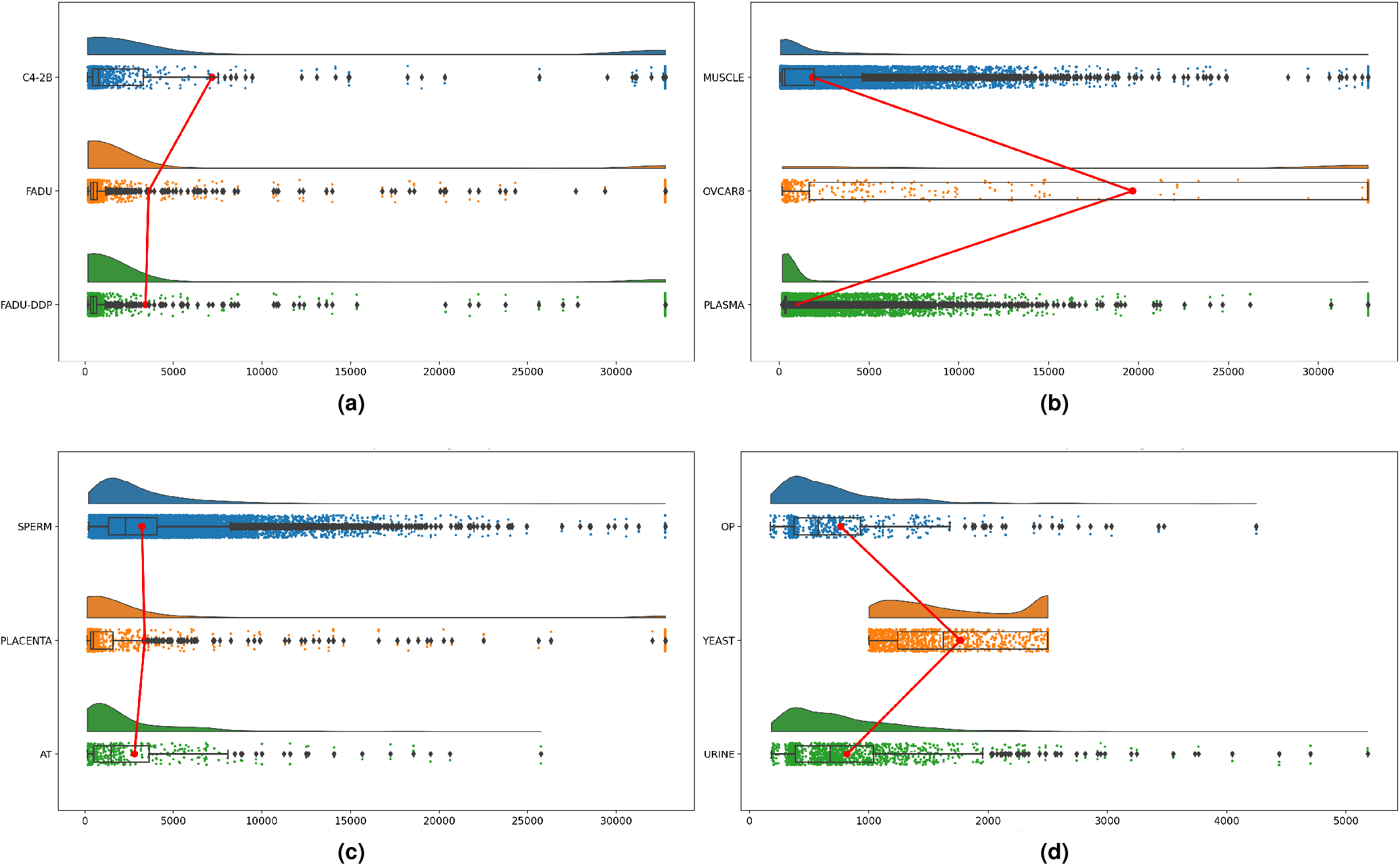
The distribution of sequence lengths across all benchmark datasets. X-axis represents the length of leccDNA and non-leccDNA sequences and y-axis represents the distribution of leccDNA and non-leccDNA sequences. The red line represents the median sequence lengths across a dataset.

#### Feature Fusion

Feature fusion involves the integration of diverse types of sequence information into a single vector which can improve the discriminative potential of statistical vectors and the efficacy of an AI predictor. Diverse types of feature fusion methods have been utilized to improve the performance of various sequence analysis tasks such as DNA hypersensitive site prediction^39^, DNA modification prediction^40^, promoter prediction^41^, and DNA binding proteins identification^42^.

In pursuit of harnessing the combined benefits of the two distinct sequence encoding methods, an early fusion strategy based on vector concatenation is adopted in the proposed iLEC-DNA predictor. Let 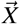 and 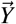 be represented as statistical vectors of dimensions P and Q for a given sequence S:

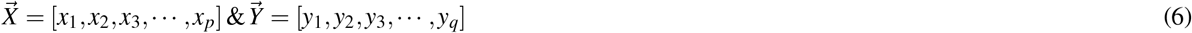

Subsequently, the fused vector can be expressed as:

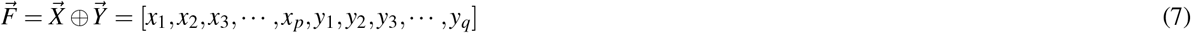

where, 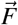 represents p+q dimensional fused vector.

### Baseline Predictors

To develop 143 baseline predictors, we utilize 13 most widely used sequence encoding methods and 11 ML classifiers. Two sequence encoding methods CKSNAP and SCPSEDNC are briefly described above. This section summarizes 11 remaining encoders namely, nucleic acid composition (NAC)^43^, enhanced nucleic acid composition (ENAC)^44^, accumulated nucleotide frequency (ANF)^45^, dinucleotide composition (DNC)^46^, trinucleotide composition (TNC)^47^, nucleotide chemical property (NCP)^48^, binary^49^, electron ionic interaction potential (EIIP)^50^, electron ionic potential values of trinucleotides (PseEIIP)^51^, pseudo dinucleotide composition (PSEDNC)^52,53^, and pseudo k-tupler composition (PSEKNC)^54^.

Nucleic acid composition (NAC)^43^ computes the normalized frequency of each nucleotide across the DNA sequence. The normalization is done through the total length of the DNA sequence. Similarly, dinucleotide composition (DNC)^46^ and trinucleotide composition (TNC)^47^, use the pairs of nucleotides (k=2, or k=3) to compute normalized occurrence frequencies rather than taking into account individual nucleotides. Enhanced nucleic acid composition (ENAC)^44^ transforms raw sequences into statistical vectors by counting the number of different k-mers at a fixed sliding window. First, a dictionary of unique k-mers is created and then for each unique each k-mer, within each window its count is computed. This step is repeated by sliding over the DNA sequences with a step size of *W*_*S*_. In the end, all the count dictionaries are concatenated together to form a discriminative statistical vector.

Accumulated nucleotide frequency (ANF)^45^ encodes nucleotide frequency information in the statistical vectors. First, it computes the position-wise counts of nucleotides and then normalizes it with the position of nucleotides. Then, it represents each nucleotide with a 4-dimensional vector at each position, where the first three values indicate the presence or absence of a specific nucleotide, and the last value is the normalized positional density of that specific nucleotide. In the binary^49^ sequence encoding method, each nucleotide is represented by a vector of size 4. These vectors include ones and zeros with each one representing the presence of a specific nucleotide.

The nucleotides of the DNA have different chemical structures and chemical properties. Phyioschemical properties-based sequence encoding methods make use of such information to capture discriminative information from the raw DNA sequences. Nucleotide chemical property (NCP)^48^, converts DNA sequences into statistical vectors based on the ring structure, functional group, and hydrogen group where each nucleotide is represented by a 3-dimensional vector. Electron-Ion Interaction Potential (EIIP)^50^ makes use of numerical values based on the average interaction potential between nucleotides constituent atoms, and electrons. It converts DNA sequences into statistical vectors by substituting each nucleotide with the predefined ionic potential value. Electron-ion interaction pseudopotentials of trinucleotide (PseEIIP)^51^ utilizes electronic ionic potential values of trinucleotides and their normalized occurrence frequency. For a trinucleotide, first the ionic potential is computed by summing up the individual pseudo-ionic potential values of three nucleotides which is multiplied by the normalized occurrence frequency of that specific trinucleotide.

Pseudo dinucleotide composition (PseDNC)^52,53^makes use of six distinct DNA properties i.e., twist, roll, rise, tilt, shift, and slide, along with the frequencies of the nucleotide pairs. First, normalized occurrence frequencies of nucleotide pairs are computed which encode the contiguous local sequence-order information of the DNA sequence. To include the global sequence-order information, a set of correlation functions are computed among the neighboring nucleotides. These functions are computed by taking the mean over the difference among the nucleotide pairs property values. The output of pseDNC is a (16+*λ*)-D vector, where the first 16 values represent the normalized frequencies of nucleotide pairs and the rest are higher-order correlation functions. Pseudo k-tupler Composition (PseKNC)^54^ works on a similar principle but the difference lies in K-tuple composition used in PseKNC. Rather than dealing only with dinucleotides or trinucleotides, PseKNC makes use of K=(1 …, L), to compute statistical vectors that contain higher and lower order features.

**Table 1.** Data statistics across cell line/tissues/isolates of AT, HM, and SC.

| HM | Samples | SC | Samples | AT | Samples |
| --- | --- | --- | --- | --- | --- |
| Muscle | 136618 | BY4741(MATa met15delta0 gene deletion) | 497 | Leaf | 167 |
| Plasma | 34953 | BY4741(MATa ura3delta0 gene deletion) | 415 | Stem | 161 |
| Sperm | 17131 | BY4741(MATa his3delta1 gene deletion) | 340 | Flower | 140 |
| FaDu | 5559 | BY4741(MATa leu2delta0 gene deletion) | 252 | Root | 73 |
| FaDu-DDP | 4539 | WT(MATa Gal2 isogenic) | 150 | AT WT Columbia-0 | 4 |
| Placenta | 3956 | WT(MATa ura3 isogenic) | 146 |  |  |
| Urine | 2182 | YCC496 | 1 |  |  |
| C4-2B | 1232 | YRH35 | 1 |  |  |
| OVCAR8 | 550 | UCC(5179, 5181) | 1 |  |  |
| Ovary-Primary | 464 | YMK(11, 23, 43, 59, 62 83, 128, 130, 132, 1453) | 1 |  |  |
| Epithelium | 14 | YJH(1069, 1305, 1310, 1315, 1372, 1414, 1462) | 1 |  |  |
| HeLa S3 | 3 |  |  |  |  |
| LAN-1, HEK-293, HeLa | 2 |  |  |  |  |
| HEK293T, HMF, F-89, SW620 | 1 |  |  |  |  |
| **Total:** | 207,211 |  | 1800 |  | 545 |

### Classifiers

This section summarizes 11 different ML classifiers that are employed along with sequence encoding methods.

Naive Bayes (NB)^55^ is a probabilistic/Bayesian model that assigns class labels to input features based on conditional probabilities of events. Based on Bayesian principles, the independence between input feature pairs is taken into account and the output of NB is the class with maximum likelihood. Logistic regression (LR)^56^ models the probability of an event by taking the log odds of an event as a linear combination of one or more independent variables. By applying transformations to linear equations, LR maps input variables to desired probability outcomes.

K-nearest neighbor (KNN)^57^ classifier works on the principle of proximity, where the data points closely related to each other are grouped together. First, the number of neighbors (k) are defined, and then a distance is computed such as Euclidean distance, or Hamming distance, and based on the distances a specific class is assigned to a data point. After computing the distances, KNN assigns new data points to the most frequently occurring class among the K nearest neighbors. This decision is based on a majority vote of adjacent data points and if K=1, data points are assigned the nearest neighbor class.

Support vector machine (SVM)^58^ creates a hyperplane that separates two classes by maximizing the distance between the margins which are the two support vectors near the decision boundary. For non-linear problems, SVM uses kernel trick to allow inner product in the mapping functions rather than the data points. Linear discriminant analysis (LDA)^59^ projects the input data into lower dimensional space such that a linear combination of features maximizes the distance between the two classes and minimizes the scatter of data in each group.

The decision tree (DT)^60^ classifier utilizes a set of rules to transform data in a tree-based structure. Initially, a node is selected based on the low Gini impurity (GI) or higher information gain (IG). After the selection of the root node, the input features are used to create further nodes in the tree based on GI scores. At each branch of the tree, a decision is made and this process is repeated recursively which also determines the next branch to follow for the final decision until a terminal node is reached. Similarly, random forest (RF)^61^ is comprised of multiple DTs working together in an uncorrelated manner. DTs in an RF work as an ensemble through the process of bootstrapping (bagging), where a class with a maximum vote from the pool of trees is chosen. Particularly, in RF each DT receives a randomly selected set of input features and samples which ensures a degree of uncorrelated decision from various DTs. In extremely randomized trees (ET)^62^, randomness is removed across the samples of the data and is induced more at the features which helps to increase the model’s accuracy as compared to RF classifier.

Adaptive boosting (AB)^63^ works by combining multiple weak classifiers which can be either decision stumps (not fully grown trees) or any other ML classifier. Initially, it passes all the samples based on individual features from the classifier and then tries to assign weights to the samples that are incorrectly classified in the process. This process is repeated multiple times until the samples are not classified correctly. Gradient boosting (GB)^64^ makes use of residual/gradients to update the weights of misclassified samples with an aim to reduce the value of these residuals such that each learner performs well in the classification of the input data.

### Benchmark Datasets

In the pursuit of creating effective and reliable machine learning (ML) predictors for any biological sequence analysis tasks, the selection of appropriate data is a crucial task^65^. Inappropriate data can lead to the development of a biased and unreliable predictor that results in misleading insights and flawed decision-making.

EccDNA sequences are available across various databases such as eccDNA Atlas^66^, TeCD^67^, EccBase^68^, EccDB^69^, and EccDNADB^70^. Each database includes extrachromosomal DNA sequences of different species and cells. Among all databases, eccDNA Atlas^66^ offers a vast and comprehensive collection of eccDNA sequences derived from diverse organisms and experimental techniques. This extensive coverage ensures a broader representation of eccDNA diversity, enabling researchers to access a more complete picture of eccDNA characteristics across different species and experimental conditions.

To prepare leccDNA identification data, first necessary details such as specie, tissue, cell, isolate, genome version, and genomic coordinates, related to leccDNA sequences are acquired from eccDNA atlas database. Specifically, the genomic coordinates encompass information such as chromosome number, start and end positions of the leccDNA sequences. In addition, the genome versions/assemblies are downloaded from Refseq (https://www.ncbi.nlm.nih.gov/refseq/)^71^, namely araport11, S288C, and hrc38. In the subsequent step, the genomic coordinates and assemblies are utilized to retrieve relevant leccDNA sequences for diverse types of species. A summary of statistics related to obtained eccDNA sequences of 3 different species i.e., *Saccharomyces cerevisiae* (SC), *Arabidopsis thaliana* (AT), and *Homo sapiens* (HS) is presented in Table **??**.

Extracted leccDNA sequences belong to 18 different cell lines in terms of HM namely, muscle (MS), plasma (PS), sperm (SP), FaDU, FaDU-DDP (FAD), placenta (PC), urine (UR), C4-2B (CB), ovcars (OV), and ovary-primary (OP), epithelium, hela, LAN-1, hela-s3, HEK-293, HEK293T, HMF, F-89, SW620, 28 different isolates of SC such as BY4741 (4 types), WT (Gal2, ura3), YCC496, YRH35, UCC(5179, 5181), YMK(1, 23, 43, 59, 62 83, 128, 130, 132, 1453), and YJH(1069, 1305, 1310, 1315, 1372, 1414, 1462). Similarly, for AT the sequences belong to 4 different tissues namely, root, stem, leaf, and flower including a wild-type AT. Due to the requirement of ML classifiers to have comprehensive training data, data related to cell lines with more than 400 samples is retained. Specifically, based on the number of samples, 9 cell lines data is discarded for HM specie such as epithelium, hela, LAN-1, hela-s3, HEK-293, HEK293T, HMF, F-89, and SW620. As the number of samples is low in terms of SC and AT, therefore two datasets from AT and SC are formulated that include leccDNA samples from all tissues/isolates.

After the retrieval of leccDNA sequences, the leccDNA sequences may contain redundant or highly similar sequences. However, these similarities can introduce a bias when dividing the data into training and testing sets, leading to an overestimation of the model performance and the establishment of impractical benchmarks. To create reliable and comprehensive benchmark datasets, following previous studies^72–74^we apply CD-HIT (sequence similarity >60%) to positive samples, where redundant or highly similar sequences are clustered together, resulting in a representative subset that encompasses the essential sequence variations. This process helps to prevent the over-representation of certain sequences, which could introduce biases during the training process of the ML classifier.

There are multiple ways to generate negative data samples for a DNA sequence classification task i.e., selection of sequences from genomic background^75,76^, and nucleotide shuffling^76,77^. For instance, sequences are randomly sampled from different positions of a genome to get a diverse pool of negative samples that are non-overlapping to the positive samples. In addition, sometimes negative samples are clustered with positive samples using psi-cd-hit to remove closely related positive and negative samples. In spite of its usage, this method has various disadvantages i.e., compositional bias, where the distribution of nucleotides in negative samples might differ completely as compared to positive samples which may lead to biased training of the ML models. In comparison, nucleotide shuffling tackles such problems by preserving various k-mers counts. Ushuffle is designed to preserve the statistical properties and local sequence features of the input sequences while removing specific sequence motifs and patterns. Following the existing work^76^, fasta_ushuffle (k=2)(https://github.com/agordon/fasta_ushuffle) is utilized to shuffle nucleotides in positive samples to obtain suitable negative samples.

### Evaluation Measures

Following evaluation criteria of existing DNA sequence classification predictors^20^–22,24, we evaluate proposed and baseline predictors using five different evaluation measures namely, accuracy (ACC), sensitivity (SN), specificity (SP), Mathews correlation coefficeint (MCC), and area under the receiver operating curve (AUC-ROC).

Accuracy^23^ refers to the proportion of correct predictions with respect to the total predictions. Specificity or true negative rate (TNR)^20^ is the model’s ability to correctly predict the negative class samples. It is determined by dividing the number of correct negative predictions by the total number of true negatives. Sensitivity (or recall)^23^ measures the ability of the model to predict positive class samples by taking the ratio of correct positive predictions to the predictions on positive samples. MCC^78^ calculates the correlation between the model predictions and the true class, by taking into consideration true positives, true negatives, false positives, and false negatives. AUC-ROC^79^ computes the degree of separability of the model based on the true positive rate (TPR) and true negative rate (TNR) at various thresholds.

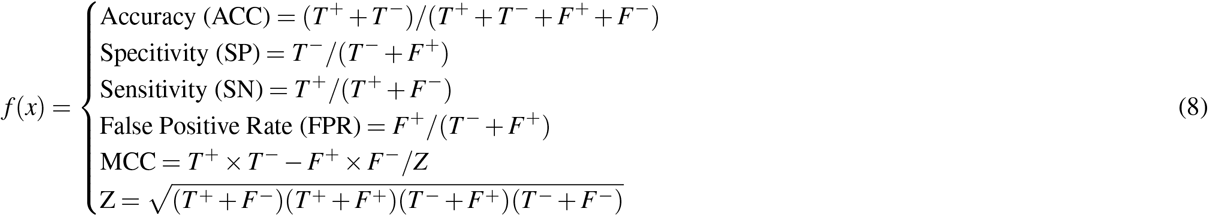

In the mathematical expression above, *T* ^+^ and *T*^*−*^ denote the true predictions related to positive and negative classes, whereas *F*^+^ and *F*^*−*^ are the incorrect predictions related to the positive and negative classes respectively.

### Experimental Setup

To prepare benchmark datasets, we utilize two different APIs namely, Biopython^80^ and USHUFFLE^77^. The proposed and baseline predictive pipelines are developed on top of two libraries namely, iLearnPlus^81^ and Scikit-Learn^82^. Following the evaluation criteria of existing DNA sequence classification predictors^20^–22,24, we perform experimentation in two different settings namely, 5-fold cross-validation and independent test set.

## Author contributions statement

A.F.A and M.N.A conceived and conducted the experiments, C.A. and D.A. analysed the results. All authors reviewed the manuscript.

## Additional information

add additional data her

